# Investigating DNA methylation as a potential mediator between pigmentation genes, pigmentary traits and skin cancer

**DOI:** 10.1101/2020.04.29.060566

**Authors:** Carolina Bonilla, Bernardo Bertoni, Josine L Min, Gibran Hemani, Genetics of DNA Methylation Consortium, Hannah R Elliott

## Abstract

**Background:** Incidence rates for melanoma and non-melanoma skin cancer (NMSC), which includes basal cell carcinoma (BCC) and squamous cell carcinoma (SCC), have been steadily increasing in all populations. Populations of European ancestry exhibit the highest rates and therefore, have been widely studied. Pigmentation characteristics are well-known risk factors for skin cancer, particularly fair skin, red hair, blue eyes and the inability to tan. Polymorphisms in established pigmentation-related genes have been associated with these traits and with an increased risk of malignancy. However, the functional relationship between genetic variation and disease is still unclear, with the exception of red hair colour variants in the melanocortin 1 receptor (*MC1R*) gene.

**Objectives:** The aim of this study was to explore the possibility that non-coding pigmentation SNPs are associated with pigmentary traits and skin cancer via DNA methylation (DNAm).

**Methods and Results:** Using a meta-GWAS of whole blood DNAm from 36 European cohorts (N=27,750; the Genetics of DNA Methylation Consortium, GoDMC), we found that 19 out of 27 pigmentation-associated SNPs distributed within 10 genes (*ASIP, BNC2, IRF4, HERC2, MC1R, OCA2, SLC24A4, SLC24A5, SLC45A2, TYR*) were associated with 391 DNAm sites across 30 genomic regions. We selected 25 DNAm sites for further analysis.

We examined the effect of the chosen DNAm sites on pigmentation traits, sun exposure phenotypes, and skin cancer, and on gene expression in whole blood. We found an association of decreased DNAm at cg07402062 with red hair in the Avon Longitudinal Study of Parents and Children (ALSPAC), and a strong positive association of DNAm at this and correlated sites with higher expression of *SPIRE2*. Additionally, we investigated the association of gene expression in skin with pigmentation traits and skin cancer. The expression of *ASIP*, *FAM83C*, *NCOA6*, *CDK10*, and *EXOC2* was associated with hair colour, whilst that of *ASIP* and *CDK10* also had an effect on melanoma and BCC.

**Conclusions:** Our results indicate that DNAm and expression of genes in the 16q24.3 and 20q11.22 regions, deserve to be further investigated as potential mediators of the relationship between genetic variants, pigmentation/sun exposure phenotypes, and some types of skin cancer.

## Introduction

The incidence of skin cancer, comprising malignant melanoma, basal cell carcinoma (BCC) and squamous cell carcinoma (SCC), has increased rapidly in the past decades[1–3]. Melanoma is the most aggressive of skin cancers although it has a low incidence, while non-melanoma skin cancer (NMSC) shows high incidence yet considerably lower mortality rates compared to melanoma[4]. To date, recognised risk factors for melanoma and NMSC include fair skin, light-coloured eyes, red hair, freckles and melanocytic naevi as well as genetic variants, some of which underlie these skin pigmentation and sun sensitivity phenotypes[5]. Even though there are missense and nonsense polymorphisms in pigmentation genes strongly associated with skin cancer, particularly in the gene *MC1R*[6], it is not fully understood how non-coding variation in these genes relates to malignancy.

DNA methylation (DNAm) is an epigenetic modification with a potential role in cancer aetiology. The association of global whole blood DNA hypomethylation with cancer is well-known and has also been described for melanoma[7,8]. More recently inter-individual DNAm variation at specific sites, measured in peripheral blood, has been uncovered as a predictor of a number of complex trait risk factors as well as all-cause mortality[9].

This study investigated whether DNAm plays a role in the relationship between genetic variants, pigmentation-related skin cancer risk factors and skin cancer. First, we examined the effect of genetic variation on DNAm levels in peripheral blood across 10 regions robustly associated with pigmentation traits and skin cancer, in 36 cohorts of European descent. We then analysed the association of pigmentation SNP-associated DNAm sites with sun exposure and pigmentation phenotypes in participants of the Avon Longitudinal Study of Parents and Children (ALSPAC). Finally, using summary data-based Mendelian randomization (SMR), we explored whether SNP-related DNAm was likely to underlie the expression of genes associated with pigmentation phenotypes and skin cancer.

## Materials and methods

The different stages of this study are depicted in Figure 1.

**Figure 1.**
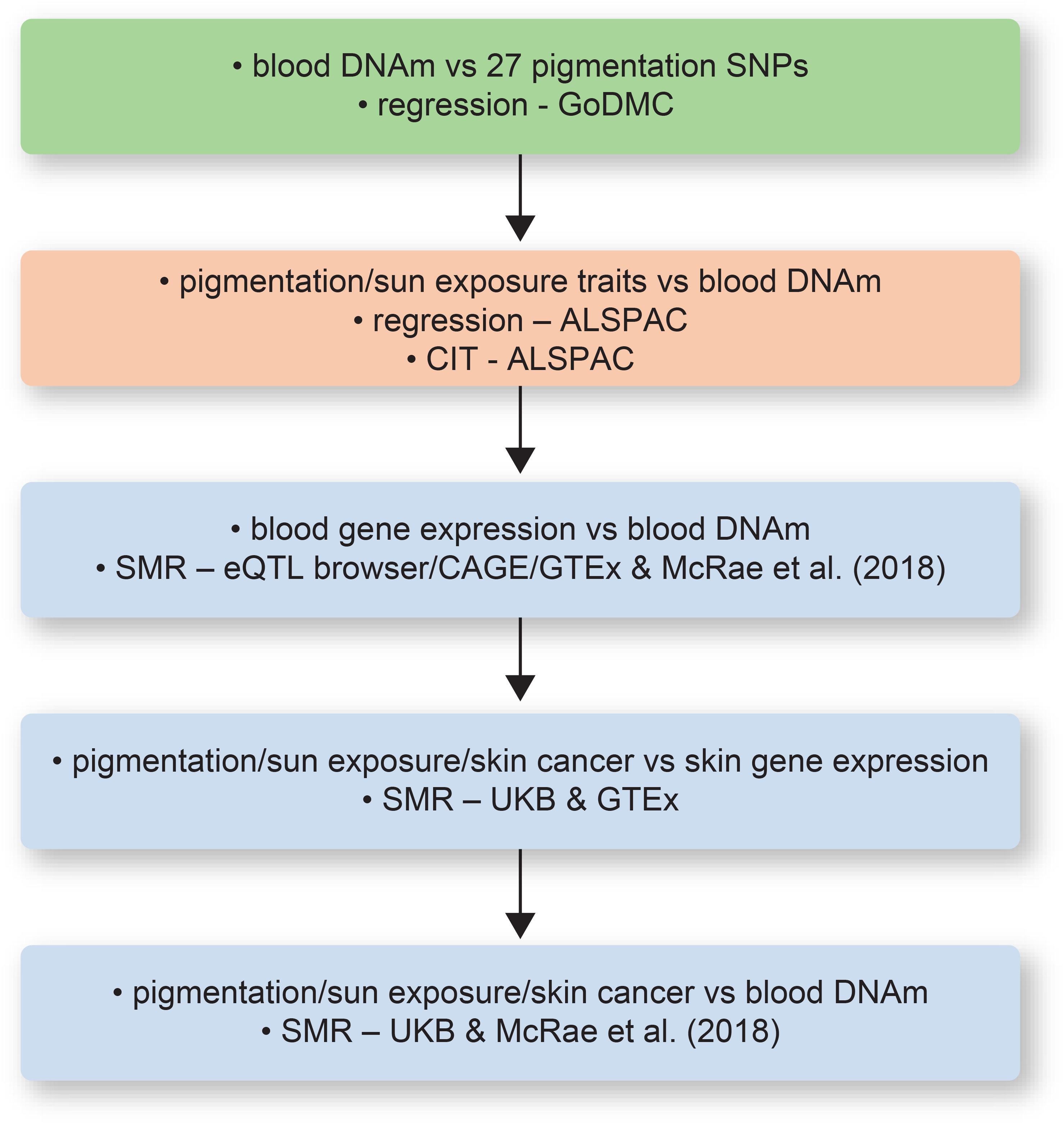
Stages of analysis implemented and datasets used in this study. From the 391 unique DNA methylation (DNAm) sites associated with pigmentation SNPs, 25 were selected for further analysis as depicted. GoDMC = Genetics of DNA methylation consortium, ALSPAC = Avon Longitudinal Study of Parents and Children, CIT = causal inference test, SMR = summary data-based Mendelian randomization, eQTL browser = blood eQTL browser, CAGE = Cap Analysis of Gene Expression, GTEx = Genotype-Tissue Expression consortium, UKB = UK Biobank. McRae et al. (2018) = reference #17.

### Identification of DNAm sites associated with pigmentation SNPs

The Genetics of DNA methylation consortium (GoDMC) was created to study the genetic basis of DNAm variation and bring together resources and researchers with expertise in the epigenetics field (www.godmc.org.uk). One of the first aims was to carry out a meta-GWAS of DNAm, as measured on Illumina 450k or EPIC Beadchips.

Results from a meta-GWAS involving 36 European cohorts (N = 27,750) were used here. Results from the GoDMC analyses are available on request from http://www.godmc.org.uk/projects.html and information on all the cohorts can be found on the GoDMC project website. We provide a brief description of the analysis in the Supplementary Methods.

Using GoDMC data, we searched for DNAm sites that were strongly associated (p < 1×10^−5^) with well-known pigmentation-related SNPs previously identified via genome-wide association studies (GWAS) or candidate gene studies (reviewed in reference [10]), located within the genes (or their surrounding regions) *ASIP* (rs1015362, rs4911414, rs619865), *BNC2* (rs2153271), *IRF4* (rs12203592, rs12210050), *HERC2* (rs12913832), *MC1R* (rs1110400, rs11547464, rs11648785, rs1805005, rs1805006, rs1805007, rs1805008, rs1805009, rs2228479, rs258322, rs4785763, rs885479), *OCA2* (rs1800401, rs1800407), *SLC24A4* (rs12896399), *SLC24A5* (rs1426654), *SLC45A2* (rs16891982, rs28777) and *TYR* (rs1042602, rs1393350).

We uncovered 874 strong SNP-DNAm associations across 30 genomic regions, which included 391 unique DNAm sites. No data was available for SNPs in *SLC24A5* and *SLC45A2*, neither for SNPs rs1110400, rs11547464, rs1805006 and rs1805009 in *MC1R*, and rs1393350 in *TYR*. Some of these polymorphisms have low minor allele frequencies (MAF) and GoDMC only included SNPs with MAF > 1%. The other SNPs may not have been part of the genotyping platforms, successfully imputed, or strongly associated with any DNAm site (p > 1×10^−5^).

Variance in DNAm explained by SNPs was estimated as 2*β^2^*MAF*(1 – MAF), where β is the effect size and MAF is the minor allele frequency.

Linkage disequilibrium (LD) r^2^ values were obtained with LDlink (https://ldlink.nci.nih.gov/).

### DNA methylation and pigmentation/sun exposure phenotypes in the Avon Longitudinal Study of Parents and Children (ALSPAC)

Cohort description and information regarding the collection of DNAm data in ALSPAC can be found in the Supplementary Methods.

#### a- regression analysis

The pigmentation/sun exposure phenotypes evaluated included: skin reflectance, freckles, moles, sunburning, tanning ability, hair colour and eye colour. We examined DNAm at the nearest time-point with respect to when the phenotypes were measured. We assessed the association of cord blood DNAm and pigmentation traits measured before age 7 [i.e. skin reflectance (49 months), freckles (49 and/or 61 months), red hair (15 and 54 months), eye colour (54 months), tanning ability (69 months)]; the association of DNAm in childhood (~7 years old) with sunburning from birth to age 12, and total number of moles at 15 years old; and the association of DNAm in adolescence (15-17 years old) with total number of moles at 15 years old, red hair at 18 years old and tanning ability at ~25 years old. Description of most of these phenotypes has been provided in an earlier work[11]. Data on hair colour and tanning ability in young adults was collected in a recent ALSPAC questionnaire (Life@25+), using the same scales employed in past questionnaires[11].

We tested the association of SNP-associated DNAm sites with pigmentation and sun exposure phenotypes using t-tests, one-way ANOVA tests, linear and logistic regression models. Regressions were adjusted for age, sex and the first 10 genetic principal components to account for population stratification. Pairwise correlation between DNAm sites was examined in ALSPAC children at age 7. All analyses were carried out with the statistical package Stata v15.

#### b- mediation analysis

The causal inference test (CIT) approach[12], implemented in the R package ‘cit’, was employed to investigate the causal direction between pigmentation SNPs, DNAm and red hair colour. Regressions were adjusted for sex, age and the top 10 genetic principal components.

### Summary data-based Mendelian randomization (SMR) of whole blood DNA methylation and gene expression

We used the platform Complex Trait Genetics Virtual Lab (CTG-VL, https://genoma.io/)[13] to run the statistical package SMR[14–16]. This program implements a method that uses summary data from GWAS, mQTL or eQTL studies to distinguish causal or pleiotropic associations between DNAm and gene expression, or between either of these and a phenotype, from a situation where the traits are caused by different variants that are in strong LD (see Figure 1b of reference [14]). If the former is true, the traits are said to be colocalized.

Peripheral blood DNAm summary data was extracted from McRae et al. [17] whilst summary data on gene expression in blood was obtained from three different sources: the Genotype-Tissue Expression consortium (GTEx, https://gtexportal.org/home/), Cap Analysis of Gene Expression (CAGE)[16], and the blood eQTL browser (https://genenetwork.nl/bloodeqtlbrowser/)[18]. All expression datasets used the hg19 genome assembly.

### Summary data-based Mendelian randomization (SMR) of gene expression in skin, pigmentation/sun exposure traits and skin cancer

We ran SMR to assess the association of gene expression in sun exposed and unexposed skin, obtained from GTEx, with pigmentary and skin cancer phenotypes.

Data was obtained by CTG-VL from the UK Biobank and trait definitions can be found in the biobank website (https://www.ukbiobank.ac.uk/). Pigmentation characteristics analysed were skin colour (UKB ID#1717, N=356,530), ease of skin tanning (#1727, N=353,697), childhood sunburn occasions (#1737, N=269,734), and black (N=15,809) and blonde (N=41,178) hair colour (#1747, N=360,270). We also considered diagnosed malignant melanoma (#ICD10:C43, N=1,672, total N=361,194), self-reported malignant melanoma (N=2,898), self-reported basal cell carcinoma (N=3,441), self-reported squamous cell carcinoma (N=449) (all #20001, total N=361,141), and having melanocytic naevi (#ICD10:D22, N=3501 cases, total N=361,194).

### Summary data-based Mendelian randomization (SMR) of whole blood DNA methylation, pigmentation/sun exposure traits and skin cancer

We also investigated the potential colocalization of genetic variants underlying DNAm with pigmentation traits and skin cancer. DNAm utilised in this analysis was measured in blood[17].

Unless otherwise reported, the SMR analyses p-value thresholds were: p SMR < 5×10^−7^/5×10^−6^ and p HEIDI ≥ 0.05.

### Heritability of the DNA methylation sites associated with pigmentation SNPs

We checked the heritability of DNAm sites using the resource provided by the Complex Disease Epigenetics Group (www.epigenomicslab.com/online-data-resources/)[19], that employs twin data to report the variance in whole blood DNAm explained by an additive genetic component, a shared environmental component and a unique environmental component.

## Results

### Identification of DNA methylation sites associated with pigmentation SNPs

We summarised available functional information on the analysed SNPs in Table 1.

**Table 1.** Pigmentation SNPs investigated in GoDMC in association with DNA methylation (DNAm) changes.

| SNP | chromosome | position | gene<sup>1</sup> | effect allele<sup>2</sup> | other allele | EAF<sup>3</sup> | DNAm sites associated | other information<sup>4</sup> |
| --- | --- | --- | --- | --- | --- | --- | --- | --- |
| rs12203592 | 6p25.3 | 396321 | *IRF4* | T | C | 0.188 | 3 | eQTL (sun exposed skin) |
| rs12210050 | 6p25.3 | 475489 | *IRF4* | T | C | 0.173 | 7 |  |
| rs2153271 | 9p22.2 | 16864521 | *BNC2* | T | C | 0.592 | 4 | eQTL (whole blood) |
| rs1042602 | 11q14.3 | 89011046 | *TYR* | A | C | 0.262 | 5 | NP_000363.1:p.Ser192Tyr |
| rs12896399 | 14q32.12 | 92773663 | *SLC24A4* | T | G | 0.468 | 6 |  |
| rs1800407 | 15q13.1 | 28230318 | *OCA2* | T | C | 0.081 | 1 | NP_000266.2:p.Arg419Gln/<br>NP_001287913.1:p.Arg395Gln |
| rs1800401 | 15q13.1 | 28260053 | *OCA2* | G | A | 0.946 | 2 | NP_000266.2:p.Arg305Trp |
| rs12913832 | 15q13.1 | 28365618 | *HERC2* | G | A | 0.744 | 3 | eQTL (whole blood) |
| rs258322 | 16q24.3 | 89755903 | *CDK10* | A | G | 0.118 | 98 | eQTL (sun exposed and unexposed skin and whole blood) |
| rs1805005 | 16q24.3 | 89985844 | *MC1R* | T | G | 0.121 | 26 | rhc (NP_002377.4:p.Val60Leu) |
| rs2228479 | 16q24.3 | 89985940 | *MC1R* | A | G | 0.089 | 163 | rhc (NP_002377.4:p.Val92Met)/eQTL (sun exposed and unexposed skin and whole blood) |
| rs1805007 | 16q24.3 | 89986117 | *MC1R* | T | C | 0.086 | 97 | RHC (NP_002377.4:p.Arg151Cys)/<br>eQTL (sun exposed and unexposed skin and whole blood) |
| rs1805008 | 16q24.3 | 89986144 | *MC1R* | T | C | 0.083 | 97 | RHC (NP_002377.4:p.Arg160Trp)/<br>eQTL (sun exposed and unexposed skin and whole blood) |
| rs885479 | 16q24.3 | 89986154 | *MC1R* | A | G | 0.105 | 110 | rhc (NP_002377.4:p.Arg163Gln)/<br>eQTL (sun exposed skin) |
| rs4785763 | 16q24.3 | 90066936 | *AFG3L1P* | A | C | 0.332 | 104 | eQTL/ eQTL (sun exposed and unexposed skin and whole blood) |
| rs11648785 | 16q24.3 | 90084561 | *DBNDD1* | C | T | 0.689 | 67 |  |
| rs4911414 | 20q11.22 | 32729444 | *ASIP* | T | G | 0.342 | 24 |  |
| rs1015362 | 20q11.22 | 32738612 | *ASIP* | C | T | 0.727 | 27 | eQTL (sun exposed and unexposed skin and whole blood) |
| rs619865 | 20q11.22 | 33867697 | *EIF6* | A | G | 0.109 | 29 |  |
| rs16891982 | 5p13.2 | 33951693 | *SLC45A2* | G | C | 0.938 | n/a | NP_001012527.2:p.Leu374Phe |
| rs28777 | 5p13.2 | 33958959 | *SLC45A2* | A | C | 0.956 | n/a |  |
| rs1393350 | 11q14.3 | 89011046 | *TYR* | A | G | 0.244 | n/a |  |
| rs1426654 | 15q21.1 | 48426484 | *SLC24A5* | A | G | 0.997 | n/a | NP_995322.1:p.Thr111Ala |
| rs1805006 | 16q24.3 | 89985918 | *MC1R* | A | C | 0.010 | n/a | RHC (NP_002377.4:p.Asp84Glu) |
| rs11547464 | 16q24.3 | 89986091 | *MC1R* | A | G | 0.009 | n/a | RHC (NP_002377.4:p.Arg142His) |
| rs1110400 | 16q24.3 | 89986130 | *MC1R* | C | T | 0.008 | n/a | RHC (NP_002377.4:p.Ile155Thr) |
| rs1805009 | 16q24.3 | 89986546 | *MC1R* | C | G | 0.008 | n/a | RHC (NP_002377.4:p.Asp294His) |

<sup>1</sup>In this study, for simplicity, SNPs on chromosome 16q24.3 are considered part of the *MC1R* genetic region, and SNPs on chromosome 20q11.22 are considered part of the *ASIP* genetic region.
<sup>2</sup>The effect allele is the allele associated with a lighter skin, hair and eye colour, and skin cancer susceptibility.
<sup>3</sup>Effect allele frequency from GoDMC for SNPs with associated DNAm sites. Effect allele frequency from 1000Genomes for SNPs not available in GoDMC.
<sup>4</sup>RHC = high penetrance red hair colour variant, rhc = low penetrance red hair colour variant. eQTL information obtained from the GTEx Portal v8 release.

Pigmentation SNPs that were strongly associated with DNAm sites in GoDMC are shown in Supplementary Table 1.

We selected DNAm sites with the most reliable effects for further analysis as follows: sites associated with at least 6 of the *MC1R* region SNPs, 3 of the *ASIP* region SNPs or showing the strongest association with pigmentation SNPs in the other genes tested. Additionally, chosen DNAm sites had to display consistent associations with the allele that increased fair pigmentation for a minimum of 5 out of 8 SNPs in *MC1R* and 2 out of 3 SNPs in *ASIP* (i.e. the allele increasing fair pigmentation had to always increase or always decrease DNAm at the same site for a majority of the SNPs tested in that gene). Finally, DNAm sites had to be associated with SNPs in ALSPAC participants consistently across the time-points where DNAm was measured (i.e. birth, childhood, adolescence, pregnancy, middle age). Based on these criteria we followed-up a total of 25 DNAm sites: 17 in *MC1R*, 3 in *ASIP*, and one each in *IRF4*, *BNC2*, *TYR*, *SLC24A4* and *HERC2*. DNAm sites associated with SNPs in *OCA2* were not present in the ARIES resource in ALSPAC so they were not taken forward. DNAm sites that were selected for further analysis are shown in Table 2.

**Table 2.**
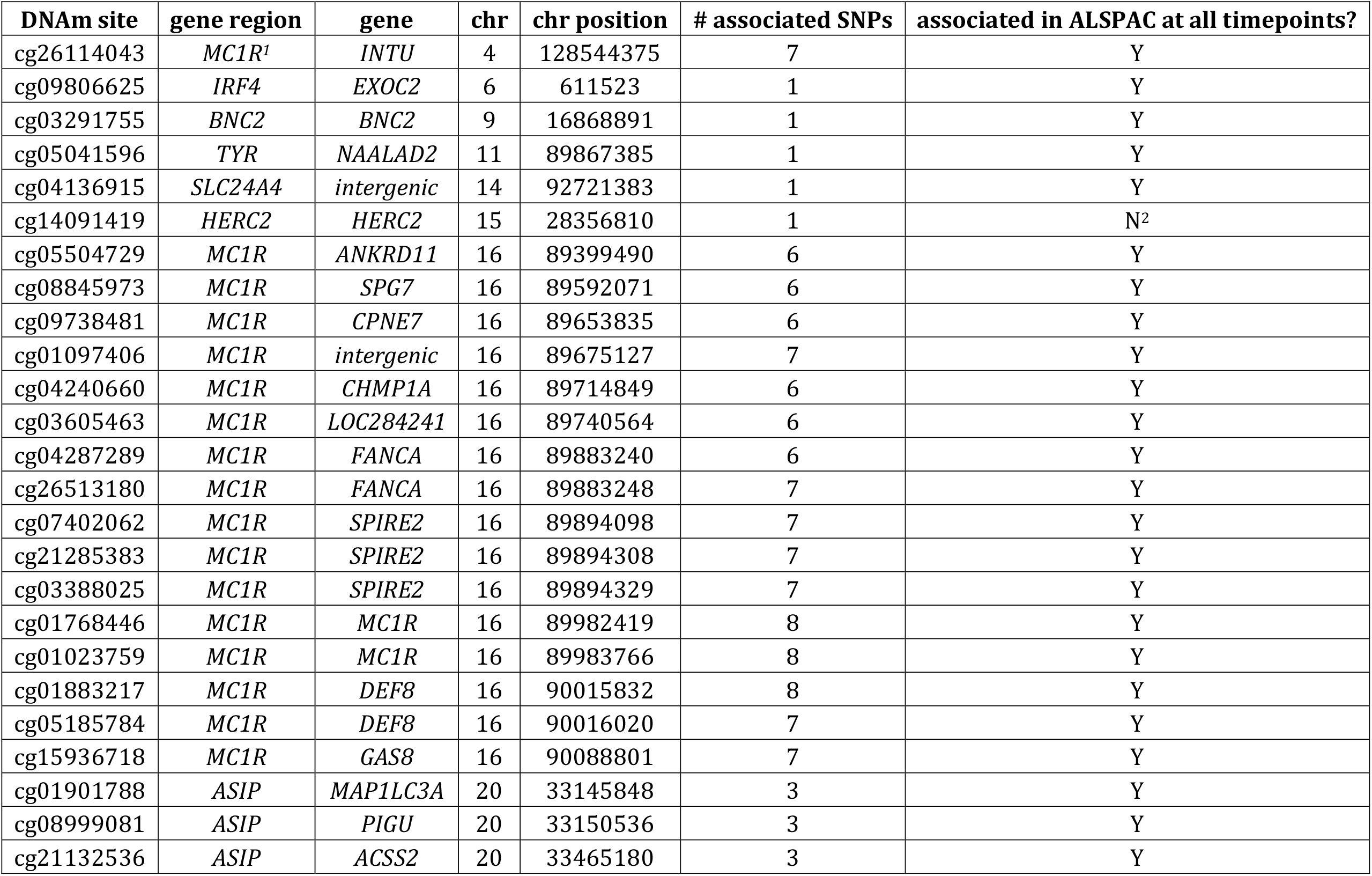
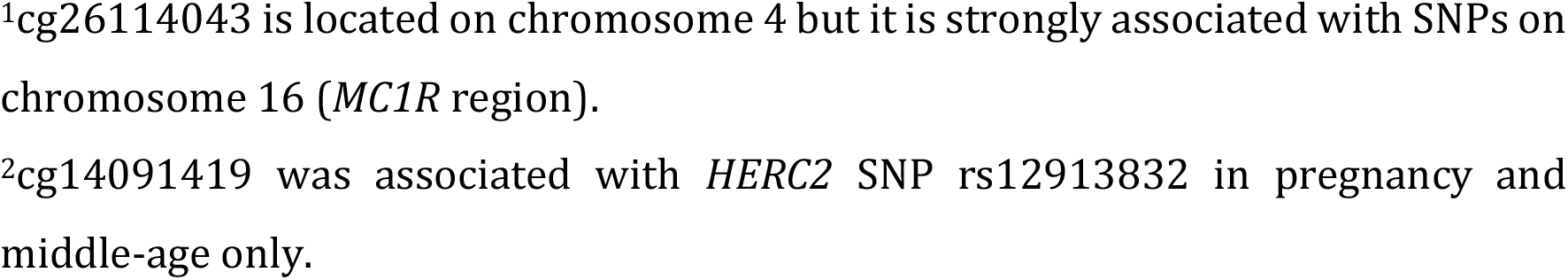
Pigmentation SNP-associated DNA methylation (DNAm) sites selected for follow-up.

| DNAm site | gene region | gene | chr | chr position | # associated SNPs | associated in ALSPAC at all timepoints? |
| --- | --- | --- | --- | --- | --- | --- |
| cg26114043 | *MC1R*<sup>1</sup> | *INTU* | 4 | 128544375 | 7 | Y |
| cg09806625 | *IRF4* | *EXOC2* | 6 | 611523 | 1 | Y |
| cg03291755 | *BNC2* | *BNC2* | 9 | 16868891 | 1 | Y |
| cg05041596 | *TYR* | *NAALAD2* | 11 | 89867385 | 1 | Y |
| cg04136915 | *SLC24A4* | *intergenic* | 14 | 92721383 | 1 | Y |
| cg14091419 | *HERC2* | *HERC2* | 15 | 28356810 | 1 | N<sup>2</sup> |
| cg05504729 | *MC1R* | *ANKRD11* | 16 | 89399490 | 6 | Y |
| cg08845973 | *MC1R* | *SPG7* | 16 | 89592071 | 6 | Y |
| cg09738481 | *MC1R* | *CPNE7* | 16 | 89653835 | 6 | Y |
| cg01097406 | *MC1R* | *intergenic* | 16 | 89675127 | 7 | Y |
| cg04240660 | *MC1R* | *CHMP1A* | 16 | 89714849 | 6 | Y |
| cg03605463 | *MC1R* | *LOC284241* | 16 | 89740564 | 6 | Y |
| cg04287289 | *MC1R* | *FANCA* | 16 | 89883240 | 6 | Y |
| cg26513180 | *MC1R* | *FANCA* | 16 | 89883248 | 7 | Y |
| cg07402062 | *MC1R* | *SPIRE2* | 16 | 89894098 | 7 | Y |
| cg21285383 | *MC1R* | *SPIRE2* | 16 | 89894308 | 7 | Y |
| cg03388025 | *MC1R* | *SPIRE2* | 16 | 89894329 | 7 | Y |
| cg01768446 | *MC1R* | *MC1R* | 16 | 89982419 | 8 | Y |
| cg01023759 | *MC1R* | *MC1R* | 16 | 89983766 | 8 | Y |
| cg01883217 | *MC1R* | *DEF8* | 16 | 90015832 | 8 | Y |
| cg05185784 | *MC1R* | *DEF8* | 16 | 90016020 | 7 | Y |
| cg15936718 | *MC1R* | *GAS8* | 16 | 90088801 | 7 | Y |
| cg01901788 | *ASIP* | *MAP1LC3A* | 20 | 33145848 | 3 | Y |
| cg08999081 | *ASIP* | *PIGU* | 20 | 33150536 | 3 | Y |
| cg21132536 | *ASIP* | *ACSS2* | 20 | 33465180 | 3 | Y |

<sup>1</sup>cg26114043 is located on chromosome 4 but it is strongly associated with SNPs on chromosome 16 (*MC1R* region).
<sup>2</sup>cg14091419 was associated with *HERC2* SNP rs12913832 in pregnancy and middle-age only.

#### MC1R

In this gene region we obtained results for red hair colour loss-of-function variants (RHC) R151C (rs1805007) and R160W (rs1805008), and missense variants (rhc) V60L (rs1805005), V92M (rs2228479) and R163Q (rs885479), and for non-coding polymorphisms found in GWAS to be associated with pigmentation and melanoma risk (rs11648785, rs258322 and rs4785763). The non-coding variants were in moderate LD with rs1805007 and between themselves (maximum r^2^ ~ 0.20), with the exception of rs258322 and rs1805007 for which r^2^ = 0.80.

*MC1R* non-coding polymorphisms and functional variants exhibited the largest number of associations with DNAm sites of all the pigmentation genes examined, ranging from 26 (rs1805005) to 163 (rs2228479) (Table 1). Three hundred and twelve unique sites were strongly associated with at least one of the *MC1R* SNPs tested. Of these, 131 were associated with just one genetic variant. There was overlap between DNAm sites associated with different *MC1R* SNPs (Supplementary Table 2). Twenty-five DNAm sites were associated with a minimum of 6 of the SNPs with results, whilst 5 were associated with all 8 of them. However, not all of these DNAm sites showed a consistent direction of association with the alleles that increase the odds of having a fair skin phenotype and being affected by melanoma. For instance, full consistency was achieved for *MC1R* DNAm site cg01097406 (i.e. all alleles that are associated with fair skin decrease DNAm at this position) and cg07130392, both associated with 7 of the 8 SNPs tested, and for cg08845973 and cg09738481, associated with 6 SNPs. Other generally consistent associations in the *MC1R* region were found for cg15936718, cg26114043 (6 agreements out of 7 associated SNPs), and cg03605463, cg04240660, and cg05504729 (5 agreements out of 6 associated SNPs).

Although most associations uncovered were between SNPs and DNAm sites in *cis*, *MC1R* showed several strong associations in *trans* (Figure 2). Besides associations with more distant DNAm sites on chromosome 16 (> 1 Mb), we identified associations with sites on chromosomes 1, 2, 3, 4, 5, 8, 10, 11, 12 and 13 (Supplementary Table 1, Figure 2). One of the *trans* sites most frequently associated with *MC1R* SNPs (7 of them, largely in low LD) was cg26114043 located in the gene *inturned planar cell polarity protein (INTU)* on chromosome 4 position 128,544,375. This gene is related to the processes of keratinocyte differentiation and hair follicle morphogenesis (https://www.ncbi.nlm.nih.gov/gene/27152).

**Figure 2.**
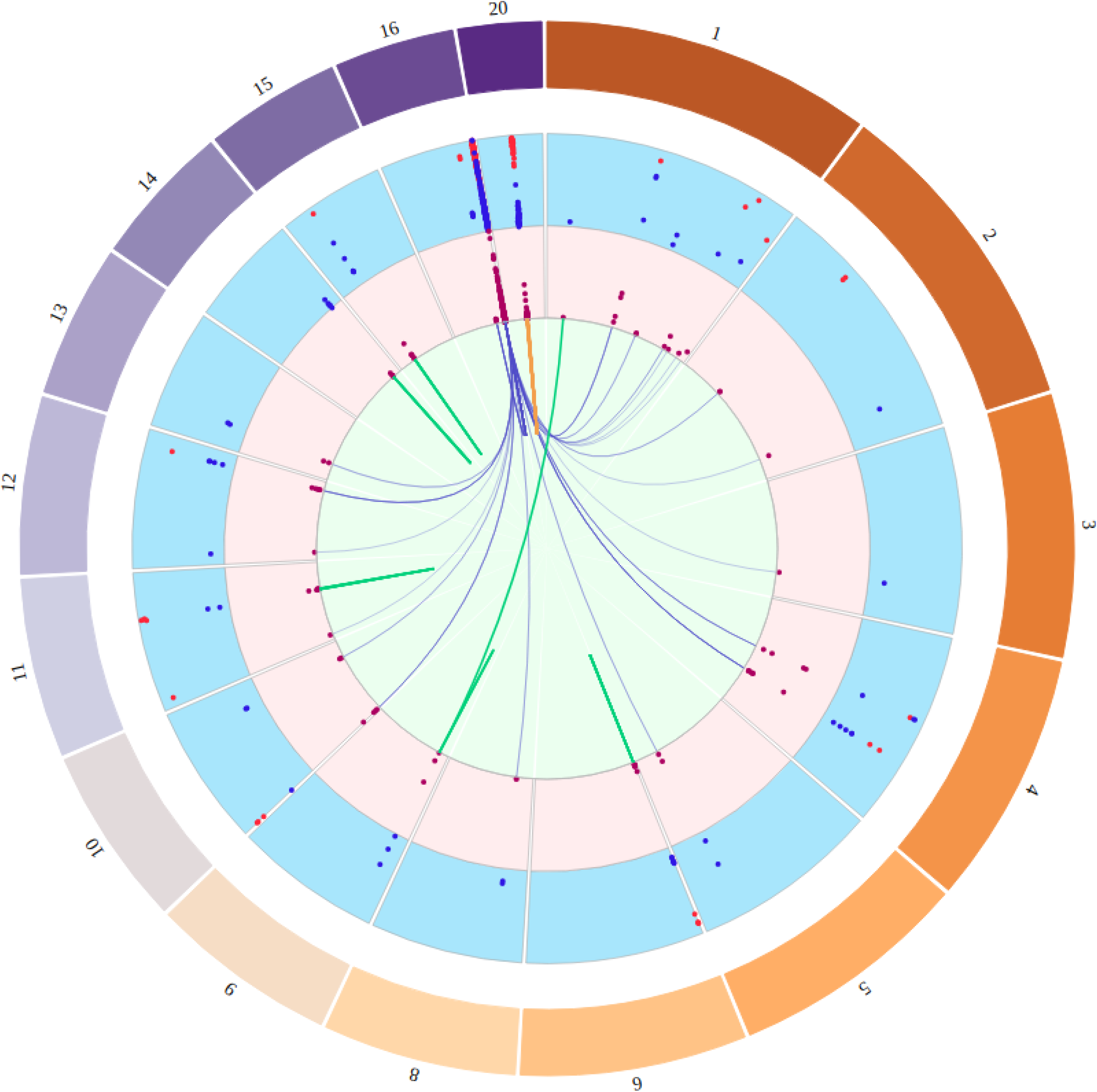
Circos plot showing the relationship between SNPs in pigmentation genes and DNA methylation (DNAm) sites in *cis* and *trans*. The external track (light blue) depicts the effect of each pigmentation SNP on DNAm (negative in red, positive in blue). The internal track (pink) depicts the R^2^ (variance explained) values. Information extracted from Supplementary Table 1.

The maximum variance in DNAm explained by a pigmentation SNP was 24.4% shown by the pair rs2228479-cg09569215. Amongst the RHC the highest R^2^ was found for rs1805007, which explained 17.8% of variation at cg08854185 (Supplementary Table 1).

#### Other pigmentation genes

The number of SNP–DNAm site associations for the other pigmentation genes examined ranged from 1 (rs1800407 in *OCA2*) to 29 (rs619865 in *ASIP*) (Table 1). LD r^2^ for variants in the same gene region was: 0.005 (rs1800401-rs1800407), 0.003 (rs1800401-rs12913832), 0.10 (rs1800407-rs12913832) in *OCA2/HERC2*; 0.04 (rs1015362-rs619865), 0.03 (rs4911414-rs619865) and 0.63 (rs1015362-rs4911414) in *ASIP*; and 0.21 (rs12203592-rs12210050) in *IRF4*.

There were 127 unique DNAm sites associated with at least one SNP in these 7 genes, whilst 102 DNAm sites were associated with only one polymorphism.

Five different DNAm sites in the *ASIP* region were associated with 3 SNPs each, whereas one site was associated with 3 SNPs in the *OCA2/HERC2* region (Supplementary Table 2). There was no DNAm site overlap between SNPs rs12210050 and rs12203592 in *IRF4*. The only fully consistent association between SNPs and DNAm sites was with cg01901788, located near *ASIP* (i.e. for all 3 SNPs the alleles associated with a fair skin phenotype increased DNAm at this site).

As expected, most associations were in *cis*, although there were three associations in *trans* in *ASIP* (distant chromosome 20) and one in *BNC2* (chromosome 1) (Supplementary Table 1, Figure 2).

Maximum variance in DNAm explained by SNPs was: 8.9% (*ASIP* rs4911414-cg08999081), 8.6% (*BNC2* rs2153271-cg03291755), 5.3% (*OCA2* rs1800407-cg20906524), 3.0% (TYR rs1042602-cg25941151), 2.7% (*IRF4* rs12210050-cg26187313), 1.7% (*HERC2* rs12913832-cg20906524), and 1.3% (*SLC24A4* rs12896399-cg16993582) (Supplementary Table 1).

### DNA methylation sites and pigmentation/sun exposure phenotypes in ALSPAC

Pairwise correlation coefficients (Pearson’s) and p-values for DNAm sites measured in ALSPAC children at age 7 are shown in Supplementary Table 3a-c. There was moderate to strong correlation between DNAm sites in the region that extends from cg04287289 to cg03388025 in *MC1R* (Supplementary Table 3b), and some moderate correlation was detected between DNAm sites in unlinked genetic regions (Supplementary Table 3a).

#### a- regression analysis

When we investigated the association of the selected DNAm sites with pigmentation and sun exposure traits in ALSPAC children we found the strongest associations with red hair colour. The most robust association with this phenotype was that of cg07402062, where redhead children showed lower levels of DNAm at this site than non-redhead children (Supplementary Table 4). This effect was seen with cord blood DNAm and red hair at 15 and 54 months (after correction for multiple testing), as well as with DNAm in adolescence and red hair at 18 years old (before correction for multiple testing). Associations with the other traits tested were either weaker or absent.

#### b- mediation analysis

The CIT showed inconclusive results for SNPs rs1805007, rs1805008, rs258322, and rs4785763 (Supplementary Table 5). There was evidence for a mediating effect of DNAm at cg07402062 between these pigmentation SNPs and red hair colour, as well as for red hair as a mediator of the relationship between the pigmentation SNPs and DNAm at cg07402062. On the other hand, for variants rs2228479 and rs885479 we found evidence of a causal effect of DNAm at cg07402062 on red hair colour, but not the other way around. Finally, rs11648785 appeared to influence both traits independently.

### Summary data-based Mendelian randomization (SMR) of whole blood DNA methylation and gene expression

Out of the 25 DNAm sites under analysis four (cg01023759, cg09738481, cg14091419, cg26114043) were not present in any of the expression datasets used. Since the data sources only included variation in *cis*, cg26114043, located on chromosome 4, was left out. The 21 remaining sites showed associations with gene expression to varying degrees which were, for the most part, consistent between datasets. Of note were the associations of correlated DNAm sites cg03388025, cg04287289, cg07402062, cg21285383 and cg26513180 with the expression of *SPIRE2*, cg08845973 with *SPG7*, cg01901788 with *NCOA6,* and cg21132536 with *GGT7*. Results of this analysis, including total numbers of associations per DNAm site and dataset, are shown in Supplementary Table 6.

### Summary data-based Mendelian randomization (SMR) of gene expression in skin, pigmentation/sun exposure traits and skin cancer

We found no evidence of a pleiotropic or causal association between gene expression at 16q24.3, 20q11.22, or any of the other pigmentation regions included in this study, and ease of skin tanning, skin colour, childhood sunburn occasions, melanocytic naevi, and SCC, in sun exposed and unexposed skin.

There was evidence of an association with the same genetic variant between black hair colour and the expression of genes *ASIP* and *FAM83C* (20q11.22) in sun exposed skin, and of *CDK10* (16q24.3), *ASIP*, *NCOA6* (20q11.22) and *EXOC2* (6p25.3) in sun unexposed skin. Whereas blonde hair colour was associated with the expression of *ASIP* and *FAM83C* in sun exposed skin, and with *NCOA6* and *EXOC2* in sun unexposed skin. In addition, *ASIP* gene expression was associated with self-reported and diagnosed malignant melanoma, and BCC in both skin tissues, and *CDK10* was associated with self-reported and diagnosed malignant melanoma, and BCC in unexposed skin (Supplementary Table 7).

Besides the regions investigated in this study, other genes displaying an association -and colocalization- were *HORMAD1*, *GOLPH3L* and *CTSS* on chromosome 1 at ~150Mb with ease of skin tanning and skin colour; and *HAL* on chromosome 12 at ~96Mb with childhood sunburn occasions. Earlier studies have reported that *HAL* participates in the initiation of the UV-B-induced immune signalling cascade, and that missense variant I439V (rs7297245) interacts with the number of severe sunburns to increase the risk for BCC and SCC[20].

When the HEIDI p-value threshold was Bonferroni-corrected for multiple testing (instead of being set as ≥ 0.05), additional genes showed potential pleiotropic/causal associations with the traits of interest, including *CPNE7*, *DBNDD1*, *DEF8*, *MC1R*, *SNAI3-AS1*, *SPATA2L*, *URAHP* in 16q24.3, and *EIF6*, *GGT7*, *UQCC*1 in 20q22.11.

We also found that skin expression of the genes *PKHD1* and *TSPAN10* previously reported as associated with hair colour[21], but which were not part of this study, was associated with ease of skin tanning, black hair, and skin colour, at the multiple comparisons corrected p-value.

### Additional summary data-based Mendelian randomization (SMR) of whole blood DNA methylation and gene expression

Because the DNAm sites we selected were associated with the expression of just 5 of the genes identified in the previous step (i.e. *DBNDD1*, *DEF8*, *GGT7*, *NCOA6*, *SPATA2L*), we searched for additional DNAm sites that showed an association with the expression of the genes reported above. For *CDK10* there was no common colocalized DNAm site across all the datasets, rather associations were evident with several DNAm sites in the 16q24.3 region. For *ASIP* and *FAM83C* there was no data available or no evidence of association with DNAm sites, and *EXOC2* showed two overlapping associated DNAm sites between the CAGE and GTEx datasets. Results for these and other genes are shown in Supplementary Table 8.

### Summary data-based Mendelian randomization (SMR) of whole blood DNA methylation, pigmentation/sun exposure traits and skin cancer

We uncovered 696 DNAm sites associated and colocalized with pigmentary and skin cancer traits (Supplementary Table 9), but of these, only a reduced number (n = 18) were associated with gene expression colocalizing with these phenotypes (see Supplementary Table 8). From this set of 18 DNAm sites, 9 were identified in our original analysis in GoDMC but only one (cg01901788) was on the list we followed up (the others were associated with 3 SNPs or less, Supplementary Table 2). SNPs associated with these “mediating” 9 DNAm sites were: rs11648785, rs1805008 (R160W), rs2228479 (V92M) and rs4785763 in the *MC1R* region; and rs1015362, rs4911414 and rs619865 in the *ASIP* region (Table 3).

**Table 3.**
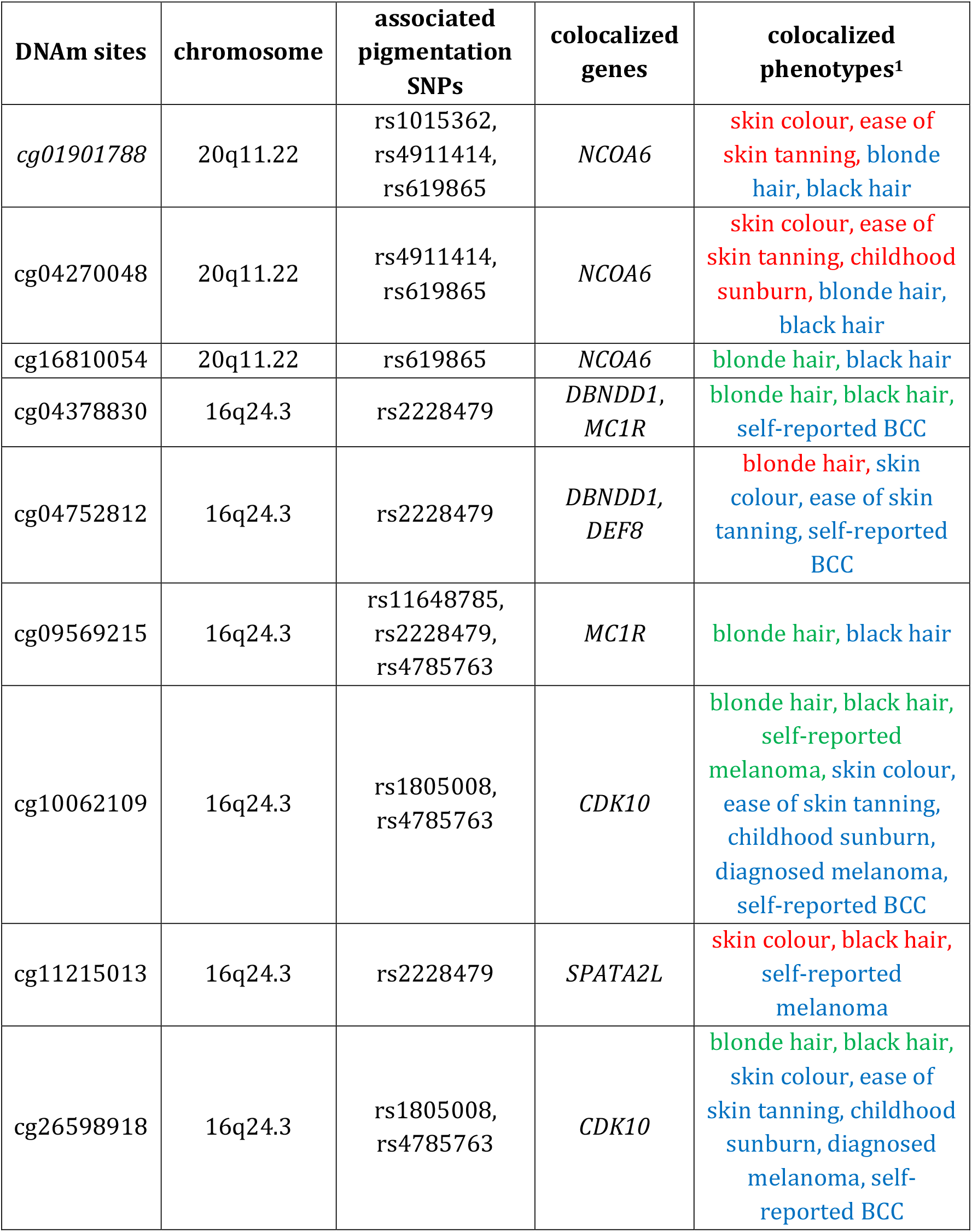
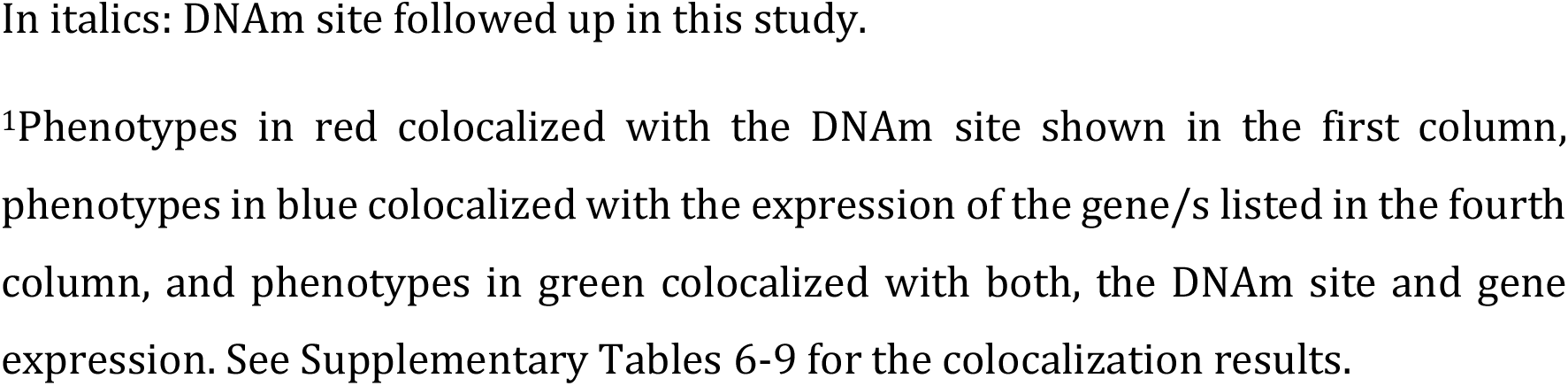
DNA methylation sites identified in GoDMC that colocalized with the expression of genes which, in turn, colocalized with pigmentation/sun exposure/skin cancer phenotypes. These DNAm sites also colocalized with pigmentation/sun exposure/skin cancer phenotypes in an independent test.

| DNAm sites | chromosome | associated pigmentation SNPs | colocalized genes | colocalized phenotypes<sup>1</sup> |
| --- | --- | --- | --- | --- |
| *cg01901788* | 20q11.22 | rs1015362,<br>rs4911414,<br>rs619865 | *NCOA6* | skin colour, ease of skin tanning, blonde hair, black hair |
| *cg04270048* | 20q11.22 | rs4911414,<br>rs619865 | *NCOA6* | skin colour, ease of skin tanning, childhood sunburn, blonde hair, black hair |
| *cg16810054* | 20q11.22 | rs619865 | *NCOA6* | blonde hair, black hair |
| *cg04378830* | 16q24.3 | rs2228479 | *DBNDD1*,<br>*MC1R* | blonde hair, black hair, self-reported BCC |
| *cg04752812* | 16q24.3 | rs2228479 | *DBNDD1*,<br>*DEF8* | blonde hair, skin colour, ease of skin tanning, self-reported BCC |
| *cg09569215* | 16q24.3 | rs11648785,<br>rs2228479,<br>rs4785763 | *MC1R* | blonde hair, black hair |
| *cg10062109* | 16q24.3 | rs1805008,<br>rs4785763 | *CDK10* | blonde hair, black hair, self-reported melanoma, skin colour, ease of skin tanning, childhood sunburn, diagnosed melanoma, self-reported BCC |
| *cg11215013* | 16q24.3 | rs2228479 | *SPATA2L* | skin colour, black hair, self-reported melanoma |
| *cg26598918* | 16q24.3 | rs1805008,<br>rs4785763 | *CDK10* | blonde hair, black hair, skin colour, ease of skin tanning, childhood sunburn, diagnosed melanoma, self-reported BCC |

In italics: DNAm site followed up in this study.
<sup>1</sup>Phenotypes in red colocalized with the DNAm site shown in the first column, phenotypes in blue colocalized with the expression of the gene/s listed in the fourth column, and phenotypes in green colocalized with both, the DNAm site and gene expression. See Supplementary Tables 6-9 for the colocalization results.

### Heritability

Nine (38%) of 24 DNAm sites followed-up showed a heritability value of 80% or more (additive genetic contribution), almost all of these are located in *MC1R*. DNAm site cg01023759 was unavailable in this dataset. Median heritability was 63% (IQR 31%, 85%) (Supplementary Table 10). The site with the largest additive genetic contribution was cg01097406 (98.6%), the site with the largest unique environmental contribution was cg09738481 (92.1%), and the site with the largest shared environmental contribution was cg05504729 (68.6%), all in *MC1R*.

## Discussion

In this study we reported the existence of strong associations between pigmentation/sun exposure/skin cancer-associated SNPs and DNAm variation at specific sites, mostly in *cis*, with a few cases in *trans*. This appears to be especially marked in the *MC1R* gene region, which could be related to the high CpG content present in this region[22]. The SNP-DNAm site associations were observed for functional variants (RHC and rhc) as well as for non-coding polymorphisms within *MC1R*.

For a set of selected DNAm sites (those associated with several pigmentation SNPs in the same direction), we assessed their effect on pigmentation/sun exposure phenotypes measured in ALSPAC participants. Although sample sizes were modest, we detected a strong inverse association of red hair with cg07402062, which is located in the gene *SPIRE2* (*MC1R* region). Interestingly, a recent GWAS of hair colour has identified the strongest genetic signal of association with red hair (vs brown/black hair) at SNP rs34357723 (T allele associated with red hair; LD with rs1805007, r^2^ = 0.27), a signal that remained relevant even after adjustment for *MC1R* coding variants[21]. Rs34357723 is an intronic variant of *SPIRE2*, and GoDMC data indicates that it is also a strong mQTL for cg07402062, explaining ~16% of DNAm at this site (T allele beta = −0.82, se = 0.01). The CIT analysis suggested that a mediation effect of cg07402062 DNAm between rs34357723 and red hair colour is compatible with ALSPAC data (see Supplementary Table 5), although replication of these findings is needed.

Galanter and colleagues reported that DNAm at cg07402062 and cg21285383 differed between ethnic groups and that these differences could be accounted for by differences in genetic ancestry[23], which would also be consistent with hair colour variation between groups.

*SPIRE2* was recently shown to participate in myosin Va-dependent transport of melanosomes in melanocytes via the generation of actin tracks[24]. However, despite *SPIRE2* expression colocalizing with DNAm at cg07402062 and correlated sites, we detected no evidence of colocalization with hair colour GWAS signals, and thus we cannot yet conclude that this gene causally underlies the phenotype.

Using summary data and two-sample Mendelian randomization implemented in SMR, we determined that the selected DNAm sites affected expression of a number of genes in *cis*, but there was limited overlap between this set of genes and those that showed an association of expression with pigmentation traits and skin cancer. Given that the DNAm-expression analysis was carried out using whole blood data, and the expression-phenotype analysis used skin data, the differences observed may be due to this discrepancy.

Previous reports on DNAm in pigmentation/skin cancer-associated regions include a study by Roos and colleagues[25] who showed that DNAm in healthy human skin was associated in *cis* with SNPs underlying melanoma risk discovered by GWAS, including two polymorphisms in *MC1R* (rs258322, rs4785763, which we tested) and one in the *ASIP* region (rs910873, not considered by us but in LD with rs619865). SNPs in *SLC45A2* (rs35390) and *TYR* (rs1393350, rs1847134) were also investigated but yielded no evidence of association with DNAm, a finding similar to ours. Roos et al. also found that none of the DNAm sites associated with these SNPs was associated with naevus count in their epigenome-wide association study (EWAS). Likewise, in our SMR analysis presence/absence of melanocytic naevi failed to colocalize with any DNAm site.

Another study by Budden and Bowden described the regulation of *MC1R* expression by a CpG island overlapping a potential enhancer in the *MC1R* gene[26]. These authors found that this CpG island is unmethylated in melanocytes but highly methylated in other skin cell types. In addition, most of the melanoma cell lines and tumours examined in their study were methylated in this region. Further analysis indicated that differential methylation of the CpG island in tumours mainly affected the expression of *MC1R*, and less so of *TUBB3*. Budden and Bowden also reported that there was weak evidence of increasing CpG island methylation levels with increasing numbers of red hair colour alleles (R) in melanoma tumours from The Cancer Genome Atlas (TCGA) dataset [26].

Whilst 7 of the 12 *MC1R* SNPs that we evaluated were located within the CpG island identified by Budden and Bowden, they were strongly associated with DNAm sites located nearby, although outside the island/enhancer region, making direct comparisons of results difficult.

Lastly, Martinez-Cadenas and colleagues[22] postulated that the considerable nucleotide diversity evident at *MC1R* was probably the result of a high mutation rate that occurred as a consequence of the elevated CpG concentration present in the region. Chromosome 16q is also one of the richest regions in gene and GC content, which have been found to be correlated[27]. In this study polymorphisms in *MC1R* were associated with DNAm variation at substantially more sites than is the case in other pigmentation genes, with lower GC content, tested (for instance, *MC1R* region SNPs are associated with 95 DNAm sites each on average, whereas *ASIP* region SNPs were associated with an average of 27 DNAm sites each, see Table 1).

From an evolutionary point of view, it is interesting to consider the potential for epigenetic modifications in pigmentation genes as a mechanism of adaptation to variable levels of ultraviolet radiation exposure, providing phenotypic flexibility in a changing environment, similar to what has been suggested for adaptation to high altitude[28]. Since there is evidence that pigmentation genes, such as *MC1R* and *ASIP*, have been shaped by natural selection[29,30], polymorphic variants in them may act, in part, by affecting how they are epigenetically regulated to optimize a response to an environment with reduced sunlight.

### Limitations

Our analysis was conducted on DNA derived from whole blood rather than skin tissue, and any pigmentation SNP-DNAm site associations may not be as strong or present in skin. We used heritability estimates provided by Hannon et al.[31] to assess the likelihood of the DNAm sites examined to vary in a similar way in different tissues. These authors reported a low contribution on average of additive genetic effects on DNAm across the genome (~16%) and a higher heritability of the sites that were correlated across tissues (median 71%). Since most of the DNAm sites (16/24) we analysed had heritability values > 50%, it is possible that the associations that we detected with pigmentation SNPs are also there in skin. Moreover, for two *MC1R* polymorphisms, Roos et al.[25] already described an association in healthy skin with some of the same DNAm sites identified in our study. Although not examining mQTLs, Yuan et al.[32] also reported an association of DNAm sites in 16q24.3 within the genes *SPG7*, *DPEP1*, *FANCA* and *SPIRE2*, which included cg04287289 and cg26513180, with ethnicity in placental tissue. However, the small overlap between the genes whose whole blood expression was affected by the DNAm sites we tested and those whose expression in skin was associated with pigmentation and skin cancer phenotypes, could be suggesting that there are in fact differences between tissues.

Assessing DNAm levels in melanocytes rather than in heterogeneous skin tissue in the future could lead to identifying new relevant sites underlying susceptibility to melanoma and NMSC[33].

Each of the methods we used have their own particular limitations, so it is advisable to be cautious in the interpretation of our findings.

The CIT method requires that the analysis is carried out in one sample where the SNP, exposure and outcome have all been measured, and therefore statistical power is constrained by the size of this sample. It may also be affected by collider bias and phenotype measurement error. In order to protect against collider bias it is essential to adjust for all confounders of the exposure-outcome relationship. Whilst we adjusted the regression models for age, sex and genetic principal components, unknown confounders may be still playing a role[34].

The SMR method was conceived to try to tell apart pleiotropy from LD in the event of an association between gene expression and a complex trait, and thus to interpret results under a model of pleiotropy rather than causality, even though it could be consistent with the latter as well[14]. To tease out a causal effect additional analyses are necessary, such as two-step Mendelian randomization[35] using independent (in *cis* or *trans*) mQTLs. However, independent instrumental variables for the DNAm sites in this study have been hard to find as there is usually considerable LD with the original SNPs. As a consequence, SMR results are mostly intended to be used to highlight genes for follow-up studies and not to establish unequivocal causal relationships[14].

Because we focused on shared DNAm sites, i.e., those that were affected by several pigmentation SNPs, and our criteria of selection were quite stringent, there were many sites with weaker associations that we did not examine in depth, which could be significant to our understanding of the function of DNAm in pigmentation genes (particularly genes besides *MC1R* and *ASIP*).

In summary, our results suggest that DNAm may lie on the causal pathway leading from pigmentation-associated SNPs to pigmentation/sun exposure traits and eventually skin cancer, possibly via regulation of gene expression, although more evidence to support these findings is needed. This study puts forward a set of genes that could be prioritised in upcoming investigations (for example, *CDK10* or *SPIRE2*) to delve deeper into the role played by variation in DNAm as a potentially causal risk factor for skin cancer.

## Acknowledgments

We are extremely grateful to all the families who took part in this study, the midwives for their help in recruiting them, and the whole ALSPAC team, which includes interviewers, computer and laboratory technicians, clerical workers, research scientists, volunteers, managers, receptionists and nurses.

## Authors’ contributions

CB conceived the study with help from BB and HRE, analysed the data and wrote the paper. HRE supervised the analysis. GH and JLM provided GoDMC data and analysis expertise. All authors critically revised the manuscript.

## Funding

CB is supported by a Universidade de São Paulo/ Coordenação de Aperfeiçoamento de Pessoal de Nível Superior fellowship (processo no. 88887.160006/2017-00).

JLM, GH and HRE are currently supported by the Medical Research Council Integrative Epidemiology Unit at the University of Bristol, UK (MC_UU_00011/5). GH is funded by the Wellcome Trust and Royal Society [208806/Z/17/Z].

The UK Medical Research Council and Wellcome (Grant ref: 217065/Z/19/Z) and the University of Bristol provide core support for ALSPAC. GWAS data for the children was generated by Sample Logistics and Genotyping Facilities at Wellcome Sanger Institute and LabCorp (Laboratory Corporation of America) using support from 23andMe. GWAS data for the mothers was generated with support from the Wellcome Trust (WT088806). Methylation data was generated with support from BBSRC (BBI025751/1 and BB/1025263/1) and from MRC (MC_UU_12013/1, MC-UU_12013/2 and MC_UU_12013/8). A comprehensive list of grant funding is available on the ALSPAC website (http://www.bristol.ac.uk/alspac/external/documents/grant-acknowledgements.pdf).

This publication is the work of the authors and CB will serve as guarantor for the contents of this paper.

## Supplementary Table legends

<u>Supplementary Table 1.</u> DNA methylation sites associated with pigmentation SNPs in the Genetics of DNA methylation Consortium (GoDMC).

Column labels are as follows: dbSNP = SNP rs#, gene = gene region, DNAm site = 450k cpg, Allele1 = effect allele, Allele2 = non-effect allele, Freq1 = effect allele frequency, FreqSE = standard error allele frequency, Effect = regression coefficient fixed effects meta-analysis, StdErr = standard error fixed effects meta-analysis, Pvalue = p-value fixed effects meta-analysis, Direction = direction for each of 36 cohorts, HetISq = I^2^, HetChiSq = heterogeneity chi-square, HetDf = degrees of freedom, HetPVal = heterogeneity p-value, EffectARE = regression coefficient additive random effects meta-analysis, StdErrARE = standard error additive random effects meta-analysis, PvalueARE = p-value random effects meta-analysis, tausq = tau square, TotalSampleSize = sample size, snpchr = SNP chromosome, snppos = SNP position build 37, cpgchr = CpG chromosome, cpgpos = CpG position build 37, cis = cis yes/no, cis = distance between SNP-CpG <1 MB, MAF = minor allele frequency, R^2^ = fraction of variability in DNAm explained by SNP.

<u>Supplementary Table 2.</u> DNA methylation (DNAm) sites associated with 2 or more pigmentation SNPs. For DNAm sites associated with 6 or more SNPs in *MC1R*, and 3 SNPs in *ASIP* and *OCA2*/*HERC2*, associations at all timepoints in ARIES are shown in red.

<u>Supplementary Table 3.</u> Pearson correlation coefficients and p-values between DNA methylation (DNAm) sites, measured in ALSPAC children at age 7. a- all DNAm sites, b-*MC1R* region, c- *ASIP* region.

<u>Supplementary Table 4.</u> Association of DNA methylation (DNAm) at cg07402062 measured at birth and adolescence with red hair at 15 months, 54 months and 18 years old in ALSPAC children.

<u>Supplementary Table 5.</u> Causal Inference Test (CIT) to evaluate the relationship between pigmentation SNPs, DNA methylation at cg07402062 and red hair colour in ALSPAC participants. p_cit = overall p-value for the CIT p_TassocL = association of the trait with the SNP genotype (p-value), p_TassocGgvnL = association of the trait with the mediator given the genotype (p-value), p_GassocLgvnT = association of the mediator with the genotype given the trait (p-value), p_LindTgvnG = genotype independent of trait given the mediator (p-value)

<u>Supplementary Table 6.</u> Summary data-based Mendelian randomization (SMR) analysis of DNA methylation (DNAm) at the 25 selected sites and gene expression (both measured in whole blood). Up to 3 genes with a p SMR < 5×10^−6^ and p HEIDI > 0.001 are shown for each DNAm site. eQTL browser = blood eQTL browser, CAGE = Cap Analysis of Gene Expression, GTEx = Genotype-Tissue Expression consortium.

<u>Supplementary Table 7.</u> Summary data-based Mendelian randomization (SMR) analysis of gene expression in sun exposed and unexposed skin and pigmentation/sun exposure/skin cancer phenotypes. All results with a p SMR below the cut-off and a p HEIDI over a Bonferroni correction are shown. Cut-offs are specified in the table in the p_SMR and p_HEIDI columns, as they vary for each phenotype. p HEIDI > 0.05 is depicted in red.

<u>Supplementary Table 8.</u> Summary data-based Mendelian randomization (SMR) analysis of DNA methylation (DNAm) and gene expression (both measured in whole blood). Only the genes showing evidence of association and colocalization in Supplementary Table 7 (plus *SPIRE2*) were examined. DNAm sites with the strongest results per gene are shown.

<u>Supplementary Table 9.</u> Summary data-based Mendelian randomization (SMR) analysis of DNA methylation (DNAm, measured in whole blood) and pigmentation/sun exposure/skin cancer phenotypes. All results with a p SMR below the cut-off and a p HEIDI over a Bonferroni correction are shown. Cut-offs are specified in the table in the p_SMR and p_HEIDI columns, as they vary for each phenotype. p HEIDI > 0.05 is depicted in red.

<u>Supplementary Table 10.</u> Heritability estimates for 25 selected DNA methylation (DNAm) sites obtained from the Complex Disease Epigenetics Group website.

## Notes

### Competing Interest Statement

The authors have declared no competing interest.

