## Supplementary Methods for "Investigating DNA methylation as a potential mediator between pigmentation genes, pigmentary traits and skin cancer"

**Cohort information**

*Genetics of DNA Methylation Consortium (GoDMC)*

Genotype data*:* Genotype data of all autosomes and chromosome X (if available) was imputed to 1000G and above using hg19/build37. Genotype data was filtered on an info score of 0.8 and a minor allele frequency (MAF) of 0.01. Genotype data was converted to bestguess data without a probability cut-off.

DNAm data*:* DNAm was measured in whole blood or cord blood using Illumina 450k or EPIC Beadchips in at least 100 European individuals. Normalised beta values were used, preferable normalised with the R package meffil[1]. Most analysts used meffil to quality control and normalize the DNAm data using functional normalization. Protocols can be found here: <https://github.com/perishky/meffil/wiki>. A github pipeline was implemented to run the analyses locally. For the genotype data, several standard sample QC steps were performed including a sex check, removal of samples with>5% missingness, and the identification and exclusion of ethnic outliers. In datasets of ostensibly unrelated individuals, those that were found to be related (identity by state > 0.125) were excluded.

The pipeline then residualised the normalised methylation betas by replacing outliers that were 10 standard deviations from the mean (3 iterations) with the probe mean, rank transforming the normalised beta values and regressing out age, sex, predicted cell counts, predicted smoking, genetic principal components and non-genetic methylation principal components. In family-based cohorts, genetic relatedness matrices were constructed and relatedness adjusted for using the GRAMMAR approach[2]. Genomic lambdas were checked by performing a GWAS of cg07959070. These residualised methylation measurements were used in all analyses.

Association analysis*:* First, every study performed a full analysis of all candidate mQTL associations, returning only associations at a threshold of p<1x10^-5^. All candidate mQTL associations at p<1x10^-5^ were combined to create a unique ‘candidate list’ of mQTL associations. In total, 102,965,711 candidate mQTL associations in *cis* (p<1x10^-5^, SNP located within 1Mb of the methylation site) and 710,638,230 candidate mQTL associations in *trans* were identified in at least one dataset. To avoid computational burden, we included *cis* associations found in at least one dataset and *trans* associations in at least two datasets. The candidate list (n=120,212,413) was then sent back to all cohorts and the association estimates obtained for every mQTL association on the candidate list.

Meta-analyses*:* Meta analyses were run using a modified version of METAL[3] using 962 chunks. Candidate mQTL associations were meta-analysed using a fixed effects model in 36 cohorts of European origin. Effects are expressed in standard deviation (SD) units of DNAm per allele.

*Avon Longitudinal Study of Parents and Children (ALSPAC)*

The ALSPAC cohort recruited pregnant women resident in Avon, UK with expected dates of delivery between 1st April 1991 to 31st December 1992. The initial number of pregnancies enrolled was 14,541. Of these initial pregnancies, there was a total of 14,676 foetuses, resulting in 14,062 live births and 13,988 children who were alive at 1 year of age. When the oldest children were approximately 7 years of age, an attempt was made to bolster the initial sample with eligible cases who had failed to join the study originally. As a result, when considering variables collected from the age of seven onwards there are data available for more than the 14,541 pregnancies mentioned above. The total sample size for analyses using any data collected after the age of seven is therefore 15,454 pregnancies, resulting in 15,589 foetuses. Of these 14,901 were alive at 1 year of age[4–6]. Study data obtained from 2014 onwards were collected and managed using Research Electronic Data Capture (REDCap) tools hosted at the University of Bristol[7,8]. A 10% sample of the ALSPAC cohort, known as the Children in Focus (CiF) group, attended clinics at the University of Bristol at various time intervals between 4 to 61 months of age. The CiF group were chosen at random from the last 6 months of ALSPAC births (1432 families attended at least one clinic).

Please note that the study website contains details of all the data that is available through a fully searchable data dictionary and variable search tool (<http://www.bristol.ac.uk/alspac/researchers/our-data/>).

Ethical approval for the study was obtained from the ALSPAC Ethics and Law Committee and the Local Research Ethics Committees. Consent for biological samples has been collected in accordance with the Human Tissue Act (2004). Informed consent for the use of data collected via questionnaires and clinics was obtained from participants following the recommendations of the ALSPAC Ethics and Law Committee at the time.

Methylation data was generated in a subset of 1,018 mother-offspring pairs from ALSPAC as part of the Accessible Resource for Integrated Epigenomics Studies (ARIES) project, using Illumina Infinium HumanMethylation450k BeadChips as described previously[9]. Following DNA extraction, samples were bisulphite converted using the Zymo EZ DNA Methylation™ kit (Zymo, Irvine, CA, USA). Following conversion, genome-wide DNAm was measured using the Illumina Infinium HumanMethylation450 (HM450) BeadChip. The arrays were scanned using an Illumina iScan, with initial quality review using GenomeStudio. ARIES was preprocessed and normalised using the *meffil* R package[1]. ARIES consists of 5469 DNAm profiles obtained from 1022 mother-child pairs measured at five time points (three time points for children: birth, childhood and adolescence; and two for mothers: during pregnancy and at middle age). Low quality profiles were removed from further processing, and the remaining 4593 profiles were normalised using the Functional Normalization algorithm{Fortin 2014} with the top 10 control probe principal components (PCs). Full details of the preprocessing and normalization of ARIES has been described previously[1].
